# Random, *de novo* and conserved proteins: How structure and disorder predictors perform differently

**DOI:** 10.1101/2023.07.18.549582

**Authors:** Lasse Middendorf, Lars A. Eicholt

## Abstract

Understanding the emergence and structural characteristics of *de novo* and random proteins is crucial for unraveling protein evolution and designing novel enzymes. However, experimental determination of their structures remains challenging. Recent advancements in protein structure prediction, particularly with AlphaFold2 (AF2), have expanded our knowledge of protein structures, but their applicability to *de novo* and random proteins is unclear. In this study, we investigate the structural predictions and confidence scores of AF2 and protein language model (pLM)-based predictor ESMFold for *de novo*, random, and conserved proteins. We find that the structural predictions for *de novo* and random proteins differ significantly from conserved proteins. Interestingly, a positive correlation between disorder and confidence scores (pLDDT) is observed for *de novo* and random proteins, in contrast to the negative correlation observed for conserved proteins. Furthermore, the performance of structure predictors for *de novo* and random proteins is hampered by the lack of sequence identity. We also observe varying predicted disorder among different sequence length quartiles for random proteins, suggesting an influence of sequence length on disorder predictions. In conclusion, while structure predictors provide initial insights into the structural composition of *de novo* and random proteins, their accuracy and applicability to such proteins remain limited. Experimental determination of their structures is necessary for a comprehensive understanding. The positive correlation between disorder and pLDDT could imply a potential for conditional folding and transient binding interactions of *de novo* and random proteins.

## Introduction

Chemically active proteins emerging from scratch, either from random sequences or from noncoding regions of the genome were once considered impossible [Zuckerkandl, 1975, Jacob, 1977]. With the development of comparative genomics, *de novo* proteins were discovered to have arisen from formerly non-coding DNA [Goffeau et al., 1996, Levine et al., 2006, Begun et al., 2007, Vakirlis et al., 2020, Keeling et al., 2019]. Devoid of sequence homology outside their evolutionary trajectory, *de novo* proteins are considered to be distant in sequence space from evolutionarily conserved proteins and might instead resemble unevolved random proteins. However, to which extent is yet unknown [Bornberg-Bauer et al., 2021, Oss and Carvunis, 2019, Heames et al., 2023]. Notably, libraries of random sequences have been shown to exhibit chemical activities and to form secondary structural elements [Keefe and Szostak, 2001, Hecht et al., 2004, Davidson et al., 1995, Tretyachenko et al., 2017]. In comparison to conserved protein, both random and *de novo* proteins are considered to be shorter and structurally more disordered, meaning they do not form a well-defined tertiary structure [Heames et al., 2023, Wilson et al., 2017, Landry et al., 2015, Basile et al., 2017, Tretyachenko et al., 2017, Dunker et al., 2008]. Nevertheless, no structure of a *de novo* protein has yet been accurately experimentally determined due to difficulties with recombinant protein expression, purification, and stability [Eicholt et al., 2022, Lange et al., 2021, Bungard et al., 2017, Aubel et al., 2023].

Advanced machine-learning-based structure predictors, most prominently AlphaFold2 (AF2) [Jumper et al., 2021], might, at least in theory, circumvent such experimental hurdles and enable structural analysis of random and *de novo* proteins *in silico*. This could help to detect novel folds, explore sequence & structure space and provide novel templates for protein engineering. Despite its unprecedented advancement, AF2 comes with certain caveats for random and *de novo* proteins. AF2 is based on co-evolutionary data which is obtained from multiple sequence alignments (MSA). These MSAs are by definition, shallow for both *de novo* and random proteins [Jumper et al., 2021]. Additionally, the aforementioned short length and high disorder of random and *de novo* proteins pose additional obstacles to structure predictions by AF2 [Aubel et al., 2023]. The presumably abundant disordered regions are flexible in space, while predictions based on the co-evolution of residues require amino acids to be in positions in a sequence which correlate to a fixed position in structure [Lindorff-Larsen and Kragelund, 2021].

As an alignment-free alternative, protein language model (pLM) based predictors have been considered to overcome the hurdles of predicting structures of proteins lacking homology, as in the case of *de novo* and random proteins [Michaud et al., 2022, Eicholt et al., 2022]. Such pLMs learn to recognize sequence architectures in proteins and their relation to structures without the need of an MSA. This process is reminiscent of learning grammar and building whole sentences and words from just the pattern of appearance of letters [Michaud et al., 2022, Ferruz and Höcker, 2022]. A significant benefit of pLM-based predictions lies in their comparatively lower computational cost and faster processing speed when compared to AF2. However, to ensure reliable structure predictions using pLM-based programs, it is essential that the sequences in the training sets are sufficiently close in sequence space to *de novo* and random proteins [Aubel et al., 2023]. ESM-Fold, which combines the final structure module of AF2 and Evolutionary Scale Modeling (ESM-2), performs high-speed predictions and is therefore, most applicable for predictions of large datasets of proteins with limited homology [Lin et al., 2023, Ahdritz et al., 2022]. Other modern structure predictors are also based on the structure module of AF2, which transforms and refines the information from the respective neural network into a protein structures [Ahdritz et al., 2022, Chowdhury et al., 2022, Wu et al., 2022]. Being based on the structure module of AF2, all modern structure predictors provide a per residue confidence score established with AF2; pLDDT (predicted local distance difference test) [Jumper et al., 2021, Mariani et al., 2013]. To calculate the pLDDT score, structure predictors use a neural network trained on predicted structures scored with per-residue lDDT-Cαagainst ground truth structures. Only high-resolution structures between (0.1-3.0 Å) were used for this training, and no NMR structures [Jumper et al., 2021]. Structure predictors provide the pLDDT from the features of the prediction itself, such as the predicted distances and angles between residues. The pLDDT score indicates the agreement between the prediction and the consensus structure obtained from multiple models of the training set rather than any particular solved structure [Jumper et al., 2021]. A low pLDDT is considered to indicate low confidence of prediction and to correlate with higher disorder [Akdel et al., 2022, Ruff and Pappu, 2021, Piovesan et al., 2022, Bruley et al., 2022, Wilson et al., 2022] and structural heterogeneity [Del Alamo et al., 2022, Saldaño et al., 2022, Alderson et al., 2022, Bruley et al., 2023]. Accordingly, pLDDT has been used for large-scale studies of structural conformations [Akdel et al., 2022, Tesei et al., 2023, Tunyasuvunakool et al., 2021, Wilson et al., 2022, Bruley et al., 2022].

Since *de novo* and random proteins are considered to be less structured and more disordered [Bornberg-Bauer et al., 2021], not only structure predictors but also appropriate disorder predictors are essential for computational analyses of their conformations [Aubel et al., 2023]. The most widely used disorder predictor is Iupred [Erdős et al., 2021], which is based on energy estimations for each amino acid independently and within the context of their neighbouring residues. It is important to note here that such energy estimations are not based on data from disordered proteins but instead from contacts between residues in experimentally resolved globular protein structures. Consequently, Iupred provides a probability for each residue how likely it is to be disorder-promoting [Erdős et al., 2021]. According to Critical Assessment of protein Intrinsic Disorder prediction (CAID) [Necci et al., 2021], deep-learning-based flDPnn outperforms Iupred and other disorder predictions on computing time and accuracy. flDPnn has also been considered to be more appropriate for random and *de novo* proteins since it is not based on evolutionary data [Aubel et al., 2023, Liu et al., 2023]. While other studies have focused on comparing different protein structure predictors for single *de novo* proteins and IDPs [Aubel et al., 2023] or single orphan proteins in comparison to selected random proteins [Liu et al., 2023], we here focus on the correlations between different predictions, their confidence and biophysical properties. Our aim is to explore how these differ on a larger scale between *de novo*, random and conserved proteins. We conducted protein structure predictions with AF2 and ESMFold [Lin et al., 2022] and annotated secondary elements using DSSP [Kabsch and Sander, 1983]. Disorder predictions were performed with both flDPnn [Hu et al., 2021] and Iupred3 (long) [Erdős et al., 2021]. While disorder correlates negatively with pLDDT for conserved proteins and IDPs as generally assumed, we found positive correlations of disorder and α-helices with pLDDT for random and *de novo* proteins. On the contrary, pLDDT correlates negatively with predictions of β-sheets in random and *de novo protein* sequences. Additionally, we quantify that MSA depth and pLDDT for *de novo* and random proteins vary, dependent on length, as proposed in Monzon *et al*. (2022) for singletons. Shorter *de novo* and random proteins show higher pLDDT per residue with lower MSA depth. Analyzing disorder and pLDDT per amino acid type, we discovered that *de novo* and conserved proteins exhibit a similar distribution. Surprisingly, the distribution of disorder per amino acid type predicted by flDPnn is uniform in random proteins. Our findings contradict the general notion of a negative correlation between disorder and pLDDT in the case of random and *de novo* proteins.

## Materials & Methods

### Dataset curation

Initially, 6716 orphan protein sequences, and their respective evolutionary age, of *Drosophila* were obtained from Heames *et al*. (2020). Redundant sequences were removed with a bias to select the sequence from *Drosophila melanogaster* if applicable. Only sequences whose mechanisms of emergence were identified as “*denovo*” or “*denovo-intron*” in Heames *et al*. (2020) were selected for analysis. This resulted in 2510 *de novo* protein sequences. Out of these 2510 proteins, 1481 were annotated as “*denovo*”, while 1029 were described as “*denovo-intron*”. Based on their date of emergence, the *de novo* proteins were divided into young (5 mya), intermediate (5-30 mya), and old (>30 mya) proteins. The three groups comprised 2205, 110, and 195 proteins, respectively. Random sequences were generated to match the amino acid distribution and sequence length of the *de novo* set as described in Heames *et al*. (2023) resulting in 2507 unique random sequences. A set of conserved proteins with the same sequence length distribution as the *de novo* proteins were randomly selected from the combined proteome of 11 *Drosophila* species. To this end, the combined proteome was filtered to match the sequence length of a respective *de novo* protein. If no sequence matched the exact sequence length of a *de novo* protein, the sequence length was step-wise increased by one amino acid until a match was found. Duplicates and *de novo* proteins were removed, resulting in 2235 unique conserved proteins. Intrinsically disordered proteins (IDPs) were selected from the DisProt database [Quaglia et al., 2021] (Release 12/2022). Proteins were included if a predicted structure was available in the AlphaFold Protein Structure Database [Varadi et al., 2021]. In total, 2205 unique IDPs were selected. Datasets are available on zenodo https://doi.org/10.5281/zenodo.7976051.

### Structure predictions

Structural predictions were performed using AlphaFold v2.1.1 on High Performance Computing Cluster PALMA II (University of Muenster). Predictions with the highest mean pLDDT were selected. AlphaFold2 predictions of DisProt proteins and conserved *Drosophila* proteins were downloaded from the AlphaFold Protein Structure Database [Varadi et al., 2021] for initial analysis. ESM-Fold [Lin et al., 2022] predictions were performed using Google Colab with a Fujitsu M520 running overnight on an RM5F, except for DisProt proteins: ESMFold on Google Colab.

### Prediction and annotation of structural features

Predicted structures of AF2 and ESMFold were used for annotation of secondary elements with DSSP (v3.0) [Kabsch and Sander, 1983]. Respective secondary element proportion was calculated by dividing the number of residues with a DSSP-annotation of “H” for α-helices, or “E” for β-sheets, by the total number of residues of the sequence. Analogously, the secondary structure of the three datasets were predicted with SPIDER3-Single [Heffernan et al., 2018] and residues annotated with “H” and “E” were used for the calculation of the secondary element proportions. For the prediction of disordered residues, Iupred3 (long) [Erdős et al., 2021] and flDPnn (default settings) [Hu et al., 2021] were used. Residues were considered disordered if their score was >= 0.5. Disorder fraction was calculated by dividing the number of disordered residues by the total sequence length.

### MSA depth calculation

The MSA depth was calculated from the feature input file generated by AF2. For each amino acid of the input sequences the number of non-gap residues were counted.

### Data & statistical analysis

Data analysis was carried out using Python (v3.9.16) [Van Rossum and Drake, 2009], Pandas (v1.4.4) [Reback et al., 2022], NumPy (v1.23.5) [Harris et al., 2020], SciPy (v1.10.1) [Virtanen et al., 2020], scikit-posthocs (v0.7.0) [Terpilowski, 2019], and BioPython (v1.80) [Cock et al., 2009]. Graphs were created using Matplolib (v3.7.1) [Hunter, 2007] and seaborn (v0.12.2) Waskom [2021]. Statistical analyzes were performed using the “kruskal” function from the scipy.stats package [Virtanen et al., 2020] (v1.10.1), followed by Dunn test while adjusting P-values based on Holm method using “Dunn” function from scikit_posthoc library [Terpilowski, 2019] (v0.7.0). Correlations of variables were calculated using “spearmanr” or “linregress” functions in scipy.stats. Data and scripts are available on GitHub: GitHub/de-novo-structure-disorder-predictor-performance.

## Results

### Predicted structural differences between random, *de novo* and conserved proteins

Due to the notion that structure predictions of random and *de novo* proteins are less confident than for conserved proteins [Aubel et al., 2023, Liu et al., 2023], we performed AF2 predictions on a set of 2510 *de novo* proteins from the *Drosophila* clade identified in Heames *et al*. (2020), random sequences matched in amino acid frequency and length (total 2507), and finally, length-matched conserved proteins from the combined *Drosophila* proteome (total 2235). As expected, we could detect a significantly lower pLDDT for random and *de novo* proteins in comparison to conserved proteins (**Figure 1A**). Disorder predictions using flDPnn of those three sets of proteins also confirmed that random and *de novo* proteins are predicted to be more disordered than conserved proteins and are following the assumed inverse relationship of low pLDDT and high disorder (**Figure 1B**). However, *de novo* proteins were predicted to be more disordered than random sequences, despite having a higher mean pLDDT (**Figure 1A & B**). When annotating secondary elements to AF2 predictions, we could detect that both random and *de novo* proteins contain secondary elements, especially α-helices, although to a lesser extent than conserved proteins (**Figure 1C & D**). After all, these predictions of disorder and annotation of secondary elements are in accordance with experimental obeservations on random and *de novo* proteins [Tretyachenko et al., 2017, Heames et al., 2023]. When comparing the secondary elements predicted by AF2 to secondary structure predictions by SPIDER3 [Heffernan et al., 2017] it is apparent that for all sets of proteins, the prediction of β-sheets is significantly different between the two programs. Additionally, only for random proteins, the predictions of α-helices are significantly different between AF2 and SPIDER3 (**Figure S1A**).

**Figure 1:**
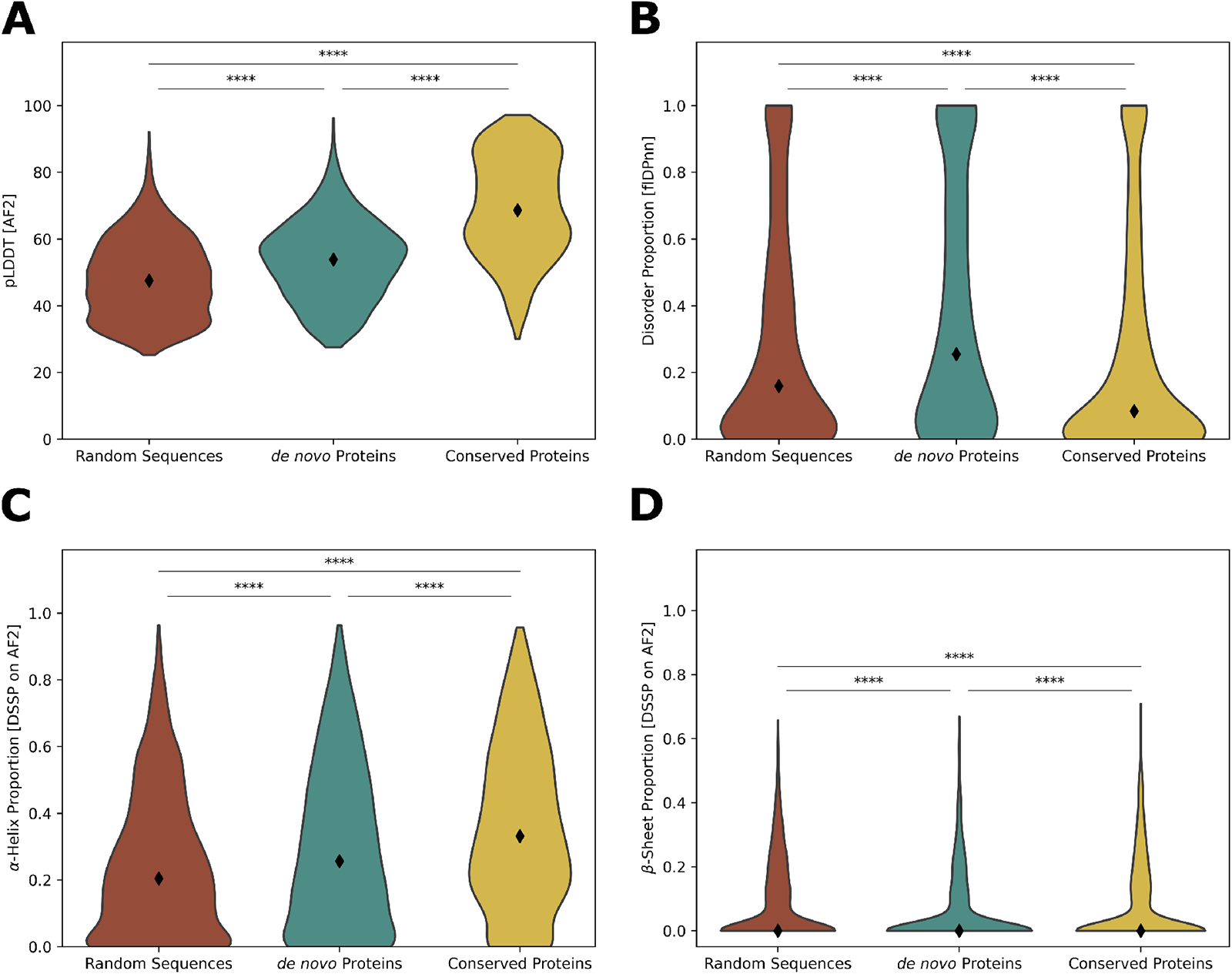
Distribution of pLDDT, disorder and DSSP annotated secondary elements for random, *de novo* and conserved proteins. **A:** Distribution of mean pLDDT from Alphafold2 predictions for random, *de novo* and conserved proteins. Mean pLDDT is higher for conserved proteins than for random and *de novo* **B:** Distribution of disorder from flDPnn. Fraction disorder of conserved proteins is predicted to be lower. **C:** Distribution of α-helices in AF2 predictions. Conserved proteins are more abundant in α-helices. **D:** Distribution of β-sheets in AF2 predictions. Significant differences are indicated by **** (P-value < 0.0001).

### For random and *de novo* proteins, pLDDT correlates negatively with **β**-sheets and positively with **α**-helices and disorder

Intrigued by the observation that AF2 predictions of *de novo* proteins have higher pLDDT values than those of random sequences despite being more disordered, we calculated the Spearman correlations of pLDDT and secondary elements (DSSP) and disorder (flDPnn) for our sets of proteins. We find that for random and *de novo* proteins, pLDDT correlated negatively with β-sheet predictions while positively for α-helices and intrinsic disorder predicted by flDPnn (**Figure 2A**). As mentioned before, it is assumed that pLDDT correlates positively with secondary elements but negatively with disorder. For conserved proteins, we could confirm this general notion of positive correlation of pLDDT with secondary elements and negative correlation with disorder (**Figure 2A**). We repeated our predictions and analysis with ESMFold and Iupred3(long) to find that the correlations persisted (**Figure 2B** & **Figure S2A & D**). Investigating correlations between residues annotated as β-sheets from AF2 and ESMFold, there is a significant difference in the case of random and *de novo* proteins, but not for conserved proteins (**Figure S2E**). Therefore, while the correlation between β-sheets and pLDDT is negative for both AF2 and ESMFold, the two programs predicted different fractions as being annotated as β-sheets (**Figure S2E**). Correlations of α-helices are near perfect between AF2 and ESMFold for all three sets of proteins (**Figure S2E**). Structural alignments between predictions of the same protein sequence by AF2 and ESMFold have low similarity in most cases, and annotations of β-sheets differ (**Figure S2B & C**). As an additional control, we performed predictions and calculated correlations for confirmed disordered proteins from DisProt database and confirmed the reported negative correlation between disorder and pLDDT (**Figure 2A**). These results indicate that the general assumption of a negative correlation between pLDDT and disorder does not apply to random and *de novo* proteins. In the case of those sets of proteins, pLDDT correlates positively with disorder and α-helices but negatively with β-sheets, both for AF2 and ESMFold.

**Figure 2:**
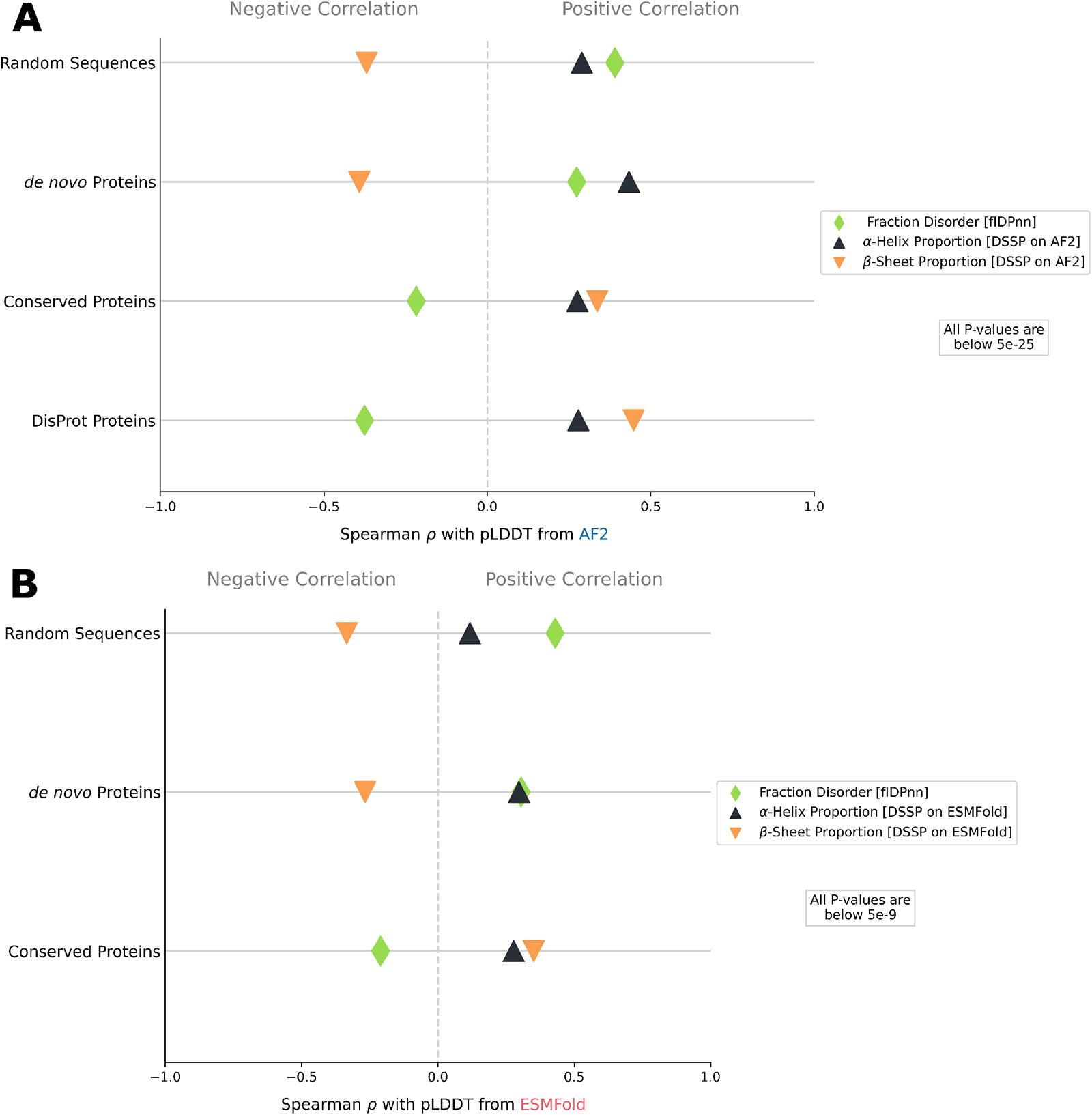
Correlation between disorder, secondary elements and pLDDT for Alphafold2 and ESMFold. **A:** Correlation between pLDDT from Alphafold2 for random, *de novo*, conserved proteins and DisProt proteins to secondary elements annotated by DSSP and disorder predicted by flDPnn. pLDDT negatively correlates with β-sheets for random and *de novo* proteins but positively for disorder and α-helices. **B:** Correlation between pLDDT from ESMFold for random, *de novo* and conserved proteins to secondary elements annotated by DSSP and disorder predicted by flDPnn. The correlations follow the same trend as for AF2.

### Influence of MSA depth & sequence length on pLDDT and disorder

Since AF2 is based on co-evolutionary information of residues extracted from multiple sequence alignments (MSA), a deeper MSA should result in a higher pLDDT [Jumper et al., 2021, Monzon et al., 2022]. When AF2 searches for sequence homology to the query, one would thus expect a deeper MSA per residue for longer sequences since these tend to align to more sequences than shorter sequences. Contrary to this, it has been noted that shorter sequences can display a higher pLDDT, despite having a lower MSA depth [Monzon et al., 2022]. Therefore, we examined correlations of MSA depth, pLDDT, and disorder to sequence length quartiles at the level of each residue (**Figure S3**). By definition, random and *de novo* proteins produce a shallow MSA, which can also be seen in the MSAs generated by AF2 (**Figure 3A & B**). As expected, the MSA depth for random sequences is lower than for *de novo* proteins. The pLDDT for random sequences decreased with increasing sequence length, while predicted disorder varies between different length quartiles, highest for the shortest sequences and lowest for the 2^nd^ length quartile (**Figure 3A**). For *de novo* proteins, we observe that, as expected, MSA depth increases with sequence length (**Figure 3B**). However, with increasing sequence length, the pLDDT decreases, despite the deeper MSA. As mentioned before, this decrease in pLDDT with increasing sequence length is, also observable for random sequences. On the other hand, with increasing sequence length, *de novo* proteins display a steady decrease in predicted disorder, unlike the varying disorder in random sequences of different lengths. For conserved proteins, the MSA depth increases with sequence length and accordingly also the pLDDT increases. In reverse, the predicted disorder decreases with increasing sequence length and indicates the negative correlation between pLDDT and disorder, independently of sequence length or MSA depth.

**Figure 3:**
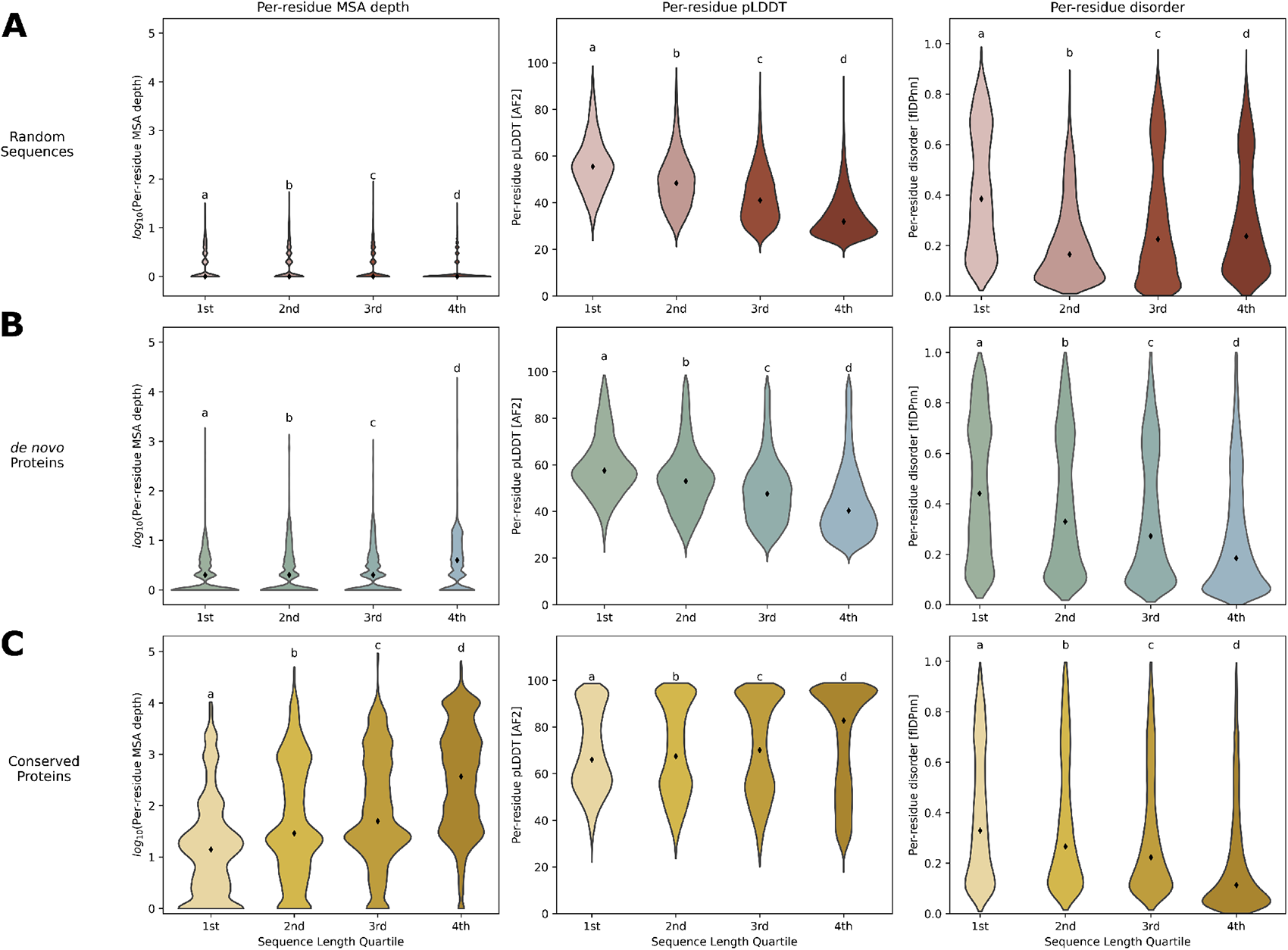
Correlations between Sequence length, MSA depth, pLDDT and disorder for random, *de novo* and conserved proteins. **A:** MSA depth, pLDDT and disorder per sequence length quartile for random sequences. 4^th^ length quartile sequences show lowest pLDDT while the 1^st^ length quartile has the highest predicted disorder. **B:** MSA depth, pLDDT and disorder per sequence length quartile for *de novo* proteins. 4^th^ length quartile sequences show the deepest MSA but lowest pLDDT and lowest disorder. **C:** MSA depth, pLDDT and disorder per sequence length quartile for conserved proteins. pLDDT and MSA depth increase with longer sequence length while disorder decreases with length. Groups annotated with different letters show statistically significant differences (P-value < 0.05).

### For random sequences, pLDDT and disorder do not change with amino acid type

Analyzing disorder values and pLDDT for each amino acid type, we observe a uniform distribution for random sequences. Primarily, the disorder values of random sequence are not influenced by the type of amino acid (**Figure 4**). On the contrary, for *de novo* and conserved proteins, both disorder and pLDDT alter depending on the type of amino acid, and their distribution is similar (**Figure 4**). As expected, more hydrophobic amino acids (C, F, I, L, V, W & Y) display lower disorder values. There is no significant difference in amino acid frequency between the *de novo* & random and the conserved set (**Figure S4**).

**Figure 4:**
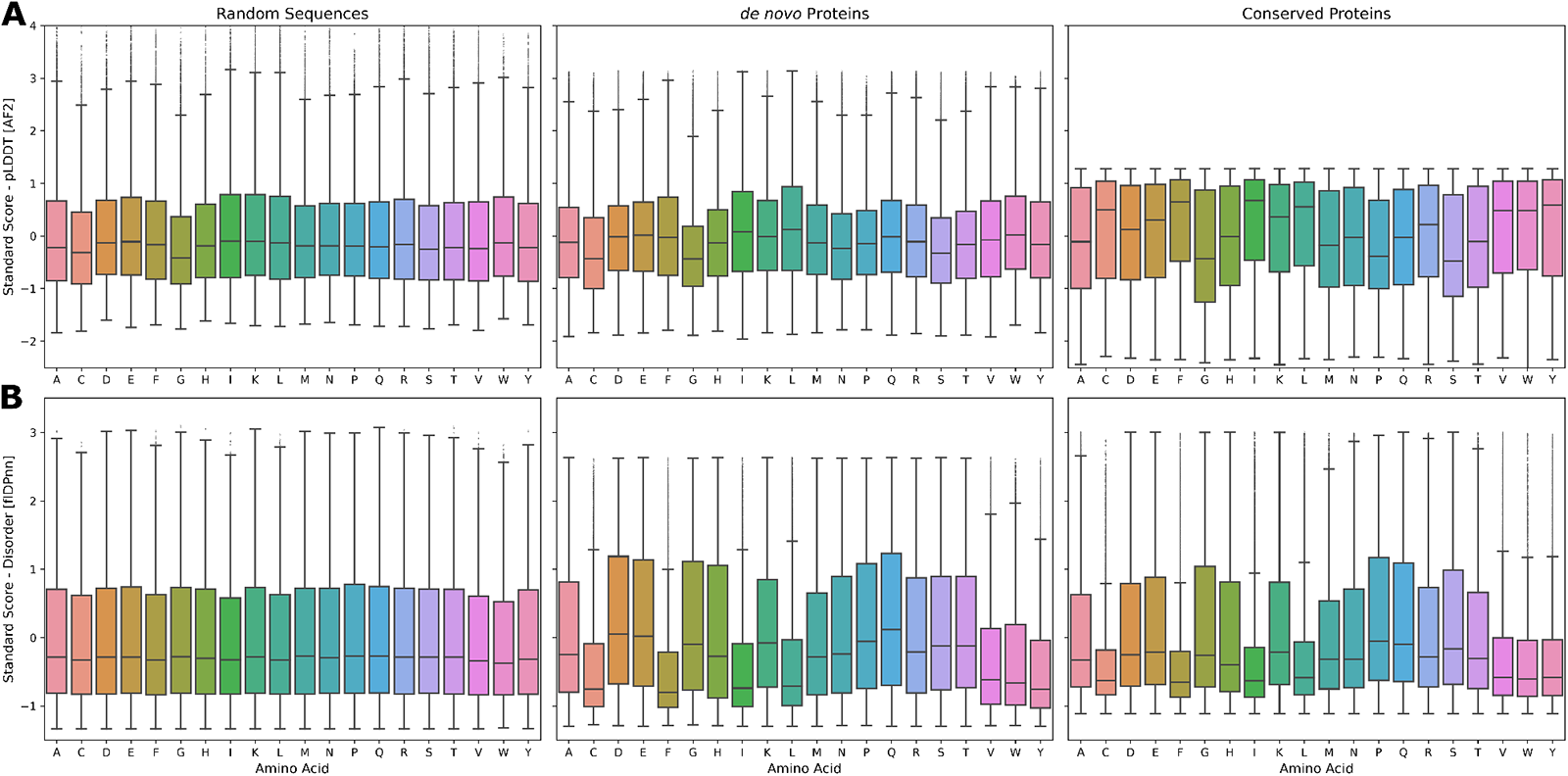
pLDDT (AF2) and disorder (flDPnn) per amino acid type in random, *de novo* and conserved proteins. **A:** pLDDT of amino acid types in random, *de novo* and conserved proteins. Differences between pLDDT per amino acid types are similar for random and *de novo* proteins but more distinct for conserved proteins. **B:** Disorder predicted by flDPnn for each amino acid type in random, *de novo* and conserved proteins. While for random sequences the distribution is quite uniform, for *de novo* and conserved proteins hydrophobic amino acids exhibit lower disorder values.

### Older *de novo* proteins are longer, have lower pLDDT and decreased disorder

Categorizing *de novo* proteins into their respective age groups (emerged 5 mya, 5-30 mya, >30 mya) [Heames et al., 2020] (**Figure 5D**), both pLDDT and disorder decrease from younger towards older *de novo* proteins (**Figure 5A**). For younger and intermediate *de novo* proteins, the disparity in pLDDT is lower than from intermediate to older *de novo* proteins. The oldest group of *de novo* proteins also contains most longer sequences (**Figure 5B & C**) and is thereby following the trend already seen for increasing sequence length leading to a decrease in both pLDDT and disorder (**Figure 3B**).

**Figure 5:**
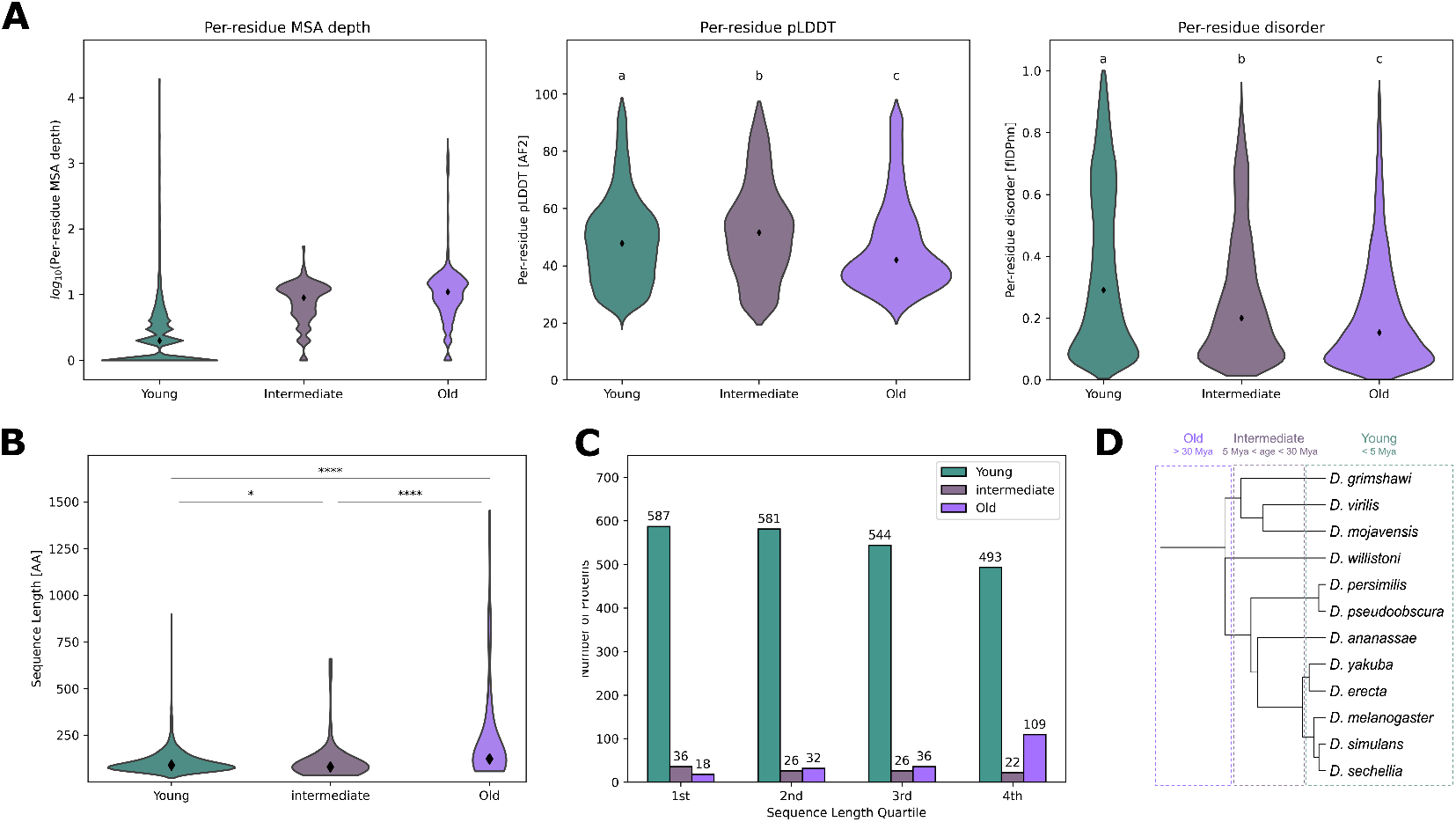
Correlations between Sequence length, MSA depth, pLDDT and disorder for different age groups of *de novo* proteins. **A:** MSA depth, per-residue pLDDT and per-residue disorder for each age group of *de novo* proteins. Older *de novo* proteins exhibit an deeper MSA, lower mean pLDDT and decrease in disorder. **B:** Mean sequence length of different age groups of *de novo* proteins. Older *de novo* proteins are longer than younger ones. **C:** Distribution of age groups per sequence length quartile. The majority of older *de novo* proteins fit into the 4^th^ quartile together with the lowest number of young sequences. **D:** Phylogenetic tree of *Drosophila* clade. Dashed boxes indicating age groups (emerged 5 mya, 5-30 mya, >30 mya). Groups annotated with different letters show statistically significant differences (P-value < 0.05). Asterisks display significant differences between groups (*: P-value < 0.05; ****: P-value < 0.0001).

## Discussion

*De novo* emerged proteins and proteins from random sequences could provide novel structures and new insights on protein structure evolution. Novel protein folds with chemical activities could contribute to a broader range of scaffolds to design new enzymes. Furthermore, proteins without ancestry, such as *de novo* and random proteins, could give us a glimpse into how the first proteins emerged. However, experimentally completely determined structures of these proteins have not been realized yet. The recent advancement of protein structure prediction has dramatically increased our knowledge of protein structures and will soon cover structure predictions of every known protein [Varadi et al., 2021, Bordin et al., 2022]. This data has already been harnessed to annotate and cluster protein structure families, identifying new folds, and to find putative *de novo* proteins [Durairaj et al., 2023, Hernandez et al., 2023]. Additionally, the pLDDT metric provided by modern structure predictors has been leveraged to evaluate structural heterogeneity and disorder [Del Alamo et al., 2022, Saldaño et al., 2022, Alderson et al., 2022, Bruley et al., 2023]. We here conclude that such kind of analyses are limited for *de novo* and random proteins, and their results from structure predictions differ highly from those of conserved proteins. We find that fractions of β-sheets and pLDDT correlate negatively for *de novo* and random proteins, instead of a negative correlation of predicted disorder and pLDDT as noted for conserved proteins (**Figure 2**) [Akdel et al., 2022, Tesei et al., 2023, Tunyasuvunakool et al., 2021]. In light of the notion, that *de novo* and random proteins are predicted to be more disordered, we repeated the analysis with experimentally confirmed disordered proteins, which displayed the same correlations as conserved proteins. Therefore, we can rule out that the positive correlation of pLDDT and disorder for *de novo* and random proteins is due to genuine, confident, ribbon-like disorder predictions by AF2. Other studies on conserved disordered proteins have found that some ribbon-like predictions by AF2, indicative of disorder, indeed exhibit a relatively high pLDDT and imply the potential of such disordered regions to undergo conditional folding and involvement in binding interactions [Piovesan et al., 2022, Brotzakis et al., 2023, Alderson et al., 2022, Zhao et al., 2023]. Such conditional folding and binding interactions could be a first step in the trajectory of *de novo* proteins to be integrated into the cellular network and become adaptive.

In line with our analysis, a recent study by Peng *et al*. (2023) further corroborates that many high-pLDDT predictions for *Drosophila de novo* proteins exhibit instability during molecular dynamics simulations under realistic conditions. This finding lends additional support to the notion that several regions characterized by high pLDDT are also predicted to be disordered. Also, the problems of AF2 with random and *de novo* sequences have been mainly attributed to a lack of sequence identity of such proteins to others. We show here that ESMFold, based on a protein language model, has the same issues as AF2, possibly due to lack of training on sequences similar to random and *de novo* proteins. Structures of random and *de novo* sequences will only be predicted with accuracy by pLM-based programs if sequences covered during training are close enough in sequence space to random and *de novo* proteins [Aubel et al., 2023]. As noted by Monzon *et al*. (2022) for proteins from AntiFam [Eberhardt et al., 2012], we also found for *de novo* and random proteins that shorter sequences result in a higher pLDDT per residue, independently of MSA depth. With decreasing pLDDT over a longer sequence length, we also see a decrease in disorder for *de novo* proteins, contradicting the inverse relationship of those two parameters. In the case of smaller proteins, the effective MSA, reflecting rare but relevant sequence identity, might lead to higher pLDDT. Also, structure predictors might be able to find the local energy minima of such small proteins while not initially being trained on biophysical data [Eicholt et al., 2022, Jumper et al., 2021]. For random sequences, we see varying predicted disorders of different sequence length quartiles. It is known that the accuracy of flDPnn varies depending on sequence length, and its most accurate for longer IDPs [Zhao et al., 2023, Necci et al., 2021]. This accuracy problem depending on sequence lengths, seems to become more vital for random proteins (**Figure 3**). Surprisingly, random proteinś pLDDT and disorder values are distributed uniformly over each amino acid type (**Figure 4**). While random sequences are not part of any program’s training set, one would assume to see differences based on the distinct biophysical properties of each amino acid. Especially since the set of random proteins matches the amino acid distribution of the *de novo* proteins. For *de novo* and conserved proteins, amino acid types resulting in higher pLDDT also give lower values of predicted disorder. The distribution of *de novo* and conserved proteins is similar, indicating that on the amino acid type level, *de novo* proteins are closer to conserved ones than to random proteins for AF2 and flDPnn. This could be due to a training set that is closer in sequence space to *de novo* proteins than to the set of random proteins generated in this study. Categorizing *de novo* proteins into three age groups, we find that older *de novo* proteins have a deeper MSA than intermediate and younger. The deeper MSA can be explained due to orthologs in several species for older *de novo* proteins, which is the basis of their age grouping. The oldest age group also displays the lowest pLDDT and lowest disorder. Once more, longer sequences have a lower pLDDT and pLDDT is not negatively correlated with disorder for *de novo* proteins.

Structure predictions can give a first glimpse at the possible structural composition of *de novo* and random proteins but it is difficult to decide which predictions are accurate. Whereas high pLDDT predictions of such proteins likely give results comparable to structural approximations [Aubel et al., 2023], we have shown here that large-scale analysis of predicted structures has many pitfalls for random and *de novo* proteins, make them a particular case for such studies. Also, the high pLDDT values in predicted disordered regions of random and *de novo* proteins could indicate their potential for conditional folding, binding and thereby, integration into the cellular network. Seeing higher disorder for younger *de novo* proteins, could imply a general evolutionary trajectory starting from disordered towards more globular structures. The combination of a dedicated disorder predictor and pLDDT scoring from structure predictors unveils high-confidence predictions pertaining to globular structures, which exhibit potential instability and significant disorder characteristics. Consequently, when dealing with random and *de novo* proteins, employment of disorder and structure predictors combined with molecular dynamics simulation, becomes imperative for comprehensive assessment of their structural properties. Whether structure predictors are reliable on *de novo* and random proteins can only be sufficiently answered with experimentally determined structures at hand [Eicholt et al., 2022, Aubel et al., 2023].

## Acknowledgements

We thank Brennen Heames, Vyacheslav Tretyachenko, and Erich Bornberg-Bauer for helpful comments on the manuscript, and Agnes Toth-Petroczy, Nobuhiko Tokuriki, Mohammed AlQuraishi and Bharat Ravi for discussing the results.

## Supporting information

Supporting figures are available on Zenodo https://doi.org/10.5281/zenodo.7976051.

## Data and Software Availability

Supplementary figures and datasets are available on Zenodo https://doi.org/10.5281/zenodo.7976051. Scripts and additional datasets are available on GitHub: GitHub/de-novo-structure-disorder-predictor-performance.

